# Learning predictive models of tissue cellular neighborhoods from cell phenotypes with graph pooling

**DOI:** 10.1101/2022.11.06.515344

**Authors:** Yuxuan Hu, Jiazhen Rong, Runzhi Xie, Yafei Xu, Jacqueline Peng, Lin Gao, Kai Tan

## Abstract

It remains poorly understood how different cell types organize and coordinate with each other to support tissue functions. We describe CytoCommunity for identification of tissue cellular neighborhoods (TCNs) based on cell phenotypes and their spatial distributions. CytoCommunity learns a mapping directly from cell phenotype space to TCN space by a graph neural network model without using additional gene or protein expression features and is thus applicable to tissue imaging data with a small number of measured features. By leveraging graph pooling, CytoCommunity enables *de novo* identification of condition-specific TCNs under the supervision of image labels. Using various types of single-cell-resolution spatial proteomics and transcriptomics images, we demonstrate that CytoCommunity can identify TCNs of variable sizes with substantial improvement over existing methods. To further evaluate the ability of CytoCommunity for discovering condition-specific TCNs by supervised learning, we apply it to colorectal and breast cancer tissue images with clinical outcome information. Our analysis reveals novel granulocyte- and cancer associated fibroblast-enriched TCNs specific to high-risk tumors as well as altered tumor-immune and tumor-stromal interactions within and between TCNs compared to low-risk tumors. CytoCommunity represents the first computational tool for end-to-end unsupervised and supervised analyses of single-cell spatial maps and enables direct discovery of conditional-specific cell-cell communication patterns across variable spatial scales.

## Introduction

To understand the structure-function relationship of a tissue, the concept of tissue cellular neighborhoods (TCNs) or spatial domains has been proposed as a recurrent functional unit in which different cell types organize and coordinate to support tissue functions ^1–3^. With the development of spatial omics and imaging technologies, there is a critical need for computational methods ^2, 4–9^ for identifying spatial domains in tissues. Several pioneering methods have been developed that can be roughly classified into statistics-based and deep learning-based approaches. As representative work in the first category, Dries *et al*. developed Giotto ^2, 4^ and Zhao *et al*. developed BayesSpace ^5^ to identify spatial domains with similar gene expression patterns based on probabilistic graphical models and spatial transcriptomics data. Chen *et al*. adapted the latent Dirichlet allocation (LDA) topic model to develop Spatial-LDA ^9^ for identifying spatially coherent patterns based on cell type counts and cell spatial coordinates. As a deep learning-based method, stLearn ^6^ utilizes a convolutional neural network model to extract features from a histology image and measures morphological similarity between neighboring cells or spots in a spatial transcriptomics image to smooth gene expression data. Clustering is then performed on the normalized expression data for spatial domain identification. Both SpaGCN ^7^ and STAGATE ^8^ first employ graph neural network to integrate gene expression and spatial location data to generate embedding representations of cells or spots in a spatial transcriptomics image and then perform clustering on those embeddings to identify spatial domains.

All existing approaches, except Spatial-LDA, are originally designed for spatial transcriptomics data and thus use expression of hundreds or thousands of genes as features to infer TCNs. On one hand, such methods may not be applicable to spatial proteomics data ^3, 10^ with single-cell resolution that only have a few tens of protein expression features available. On the other hand, using spatial transcriptomic data as inputs cannot directly establish the relationship between cell types and TCNs in a tissue, making the interpretation of TCN identity and function challenging. Furthermore, given a cohort of tissue images associated with different conditions (e.g. disease risk and patient prognosis), it is critical to identify condition-specific TCNs in order to discover TCNs with more biological and clinical relevance. A representative condition-specific TCN in cancer tissues is the tertiary lymphoid structure, which is typically present in low-risk but absent in high-risk patients of many cancer types ^11^. All existing methods are designed to detect TCNs in individual tissue images and thus not applicable to identification of condition-specific TCNs because alignment of TCNs across images is NP-hard ^12^. To our knowledge, no method currently exists for *de novo* identification of condition-specific TCNs by utilizing tissue image labels explicitly.

Here, we describe the CytoCommunity algorithm for identifying TCNs that can be applied in either an unsupervised or a supervised learning framework. We formulate the TCN identification as a graph-based community detection problem and employ a minimum-cut-based graph neural network model to identify TCNs in cell-cell spatial proximity graphs with cell types as node attributes. CytoCommunity directly uses cell types as initial features of cells to learn TCNs and thus can be applied to both single-cell transcriptomics and proteomics data and facilitates the interpretation of TCN functions. CytoCommunity can not only infer TCNs from individual images but also identify condition-specific TCNs from a cohort of labeled tissue images by leveraging graph pooling and image labels, which is an effective strategy to address the difficulty of graph alignment. Our deep graph learning framework is different from the group of methods represented by SpaGCN ^7^ and STAGATE ^8^, which employ a “cell embedding and clustering” procedure to identify spatial domains. The clustering approach makes it difficult to adopt a supervised learning framework for finding condition-specific tissue structures.

Using single-cell spatial proteomics and transcriptomics datasets, we benchmarked the performance of unsupervised CytoCommunity on detection of TCNs of variable sizes in individual healthy tissue images. We also demonstrated the ability of supervised CytoCommunity to reveal the changes in within- and between-TCN communications in tumor tissues of patients having different risks and prognoses.

## Results

### Overview of the CytoCommunity algorithm

The CytoCommunity algorithm consists of two components: a graph neural network (GNN)-based soft tissue cellular neighborhood (TCN) assignment module and a TCN ensemble module to determine the final robust TCNs in single-cell spatial maps (Fig. 1a). CytoCommunity can be used for both unsupervised (Fig. 1b; Online Methods) and supervised learning tasks (Fig. 1c; Online Methods). As an unsupervised learning task, given a single-cell spatial map with cell type and location information, CytoCommunity first constructs a k-nearest-neighbor (KNN) graph with nodes representing cells and edges representing the spatial proximity among cells. Each node also has an attribute vector representing its associated cell phenotype (e.g. cell type or state). Using a deep neural network consisting of a graph convolution layer and a fully-connected layer, the node attribute vectors on the KNN graph are transformed to the soft TCN assignment vectors, representing probabilities of cells belonging to the specified number of TCNs. A graph minimum cut (MinCut)-based loss function ^13^ is applied to learn optimal soft TCN assignments for cells. This loss function can be used alone to detect TCNs in individual single-cell spatial maps without any image labels (e.g. disease stage of the corresponding patient sample). In a supervised learning task for *de novo* identification of condition-specific TCNs, a second deep neural network with a graph pooling layer, a graph convolution layer and two fully-connected layers are added to the soft TCN assignment module. A cross-entropy loss function is used for image classification to minimize the difference between actual and predicted probabilities of image labels. The overall loss function of the supervised learning is a linear combination of the MinCut-based and the cross-entropy-based loss functions. Due to the joint training of the two loss functions, the learned optimal soft TCN assignments of cells are associated with the conditions/labels of the single-cell spatial images under study. To alleviate the instability of the graph partitioning based on GNN, the soft TCN assignment module can be run multiple times to generate multiple optimal soft TCN assignment matrices. Finally, a majority-voting-based ensemble procedure is performed on these soft TCN assignment matrices to determine the final TCNs.

**Figure 1.**
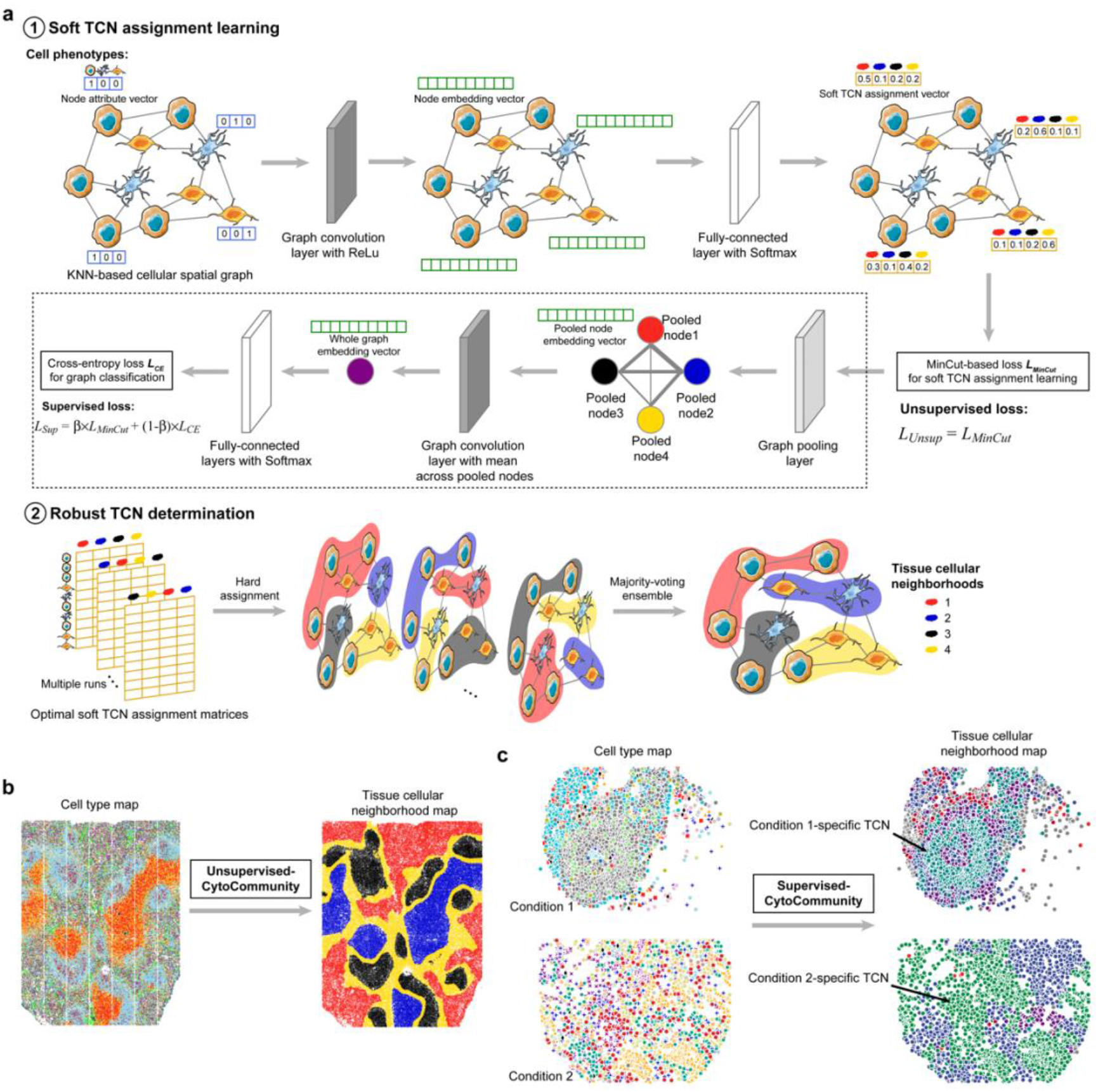
Schematic diagram of the CytoCommunity algorithm. Given a single-cell spatial map with cell type annotation and cell spatial coordinates, identification of tissue cellular neighborhoods (TCNs) is formulated as a graph partitioning problem, which can be solved by a graph neural network (GNN)-based machine learning model. **(a)** The CytoCommunity algorithm includes two components: a soft TCN assignment module and a TCN ensemble module to determine the final robust TCNs. First, an undirected K-nearest neighbor (KNN) graph is constructed based on Euclidean distance between cells computed using the cell spatial coordinates. Each node represents a cell and its attribute vector (blue) is represented using one-hot encoding. An edge exists between two nodes if a node belongs to the KNN set of the other node. A basic GNN model with the ReLu activation function is applied to this node-attributed KNN graph to obtain a real-valued embedding vector (green) for each node, which integrates the phenotype information of the cell and its local neighborhood. *d*, the number of embedding dimensions. A fully-connected neural network (no hidden layers) with the softmax activation function is used for transforming the node embeddings to the soft TCN assignments (yellow vectors) of nodes, representing the probabilities of cells belonging to the specified number of TCNs. The graph minimum cut-based loss (*L_MinCut_*) function is then used to learn the optimal soft TCN assignments of all nodes. This loss function can be used alone for an unsupervised learning task. In a supervised learning task for *de novo* identification of condition-specific TCNs in a tissue image dataset, a graph pooling, a graph convolution and two fully-connected layers with the cross-entropy loss function *L_CE_* (for image classification, surrounded by a dashed rectangular box) are added on top of the soft TCN assignment module. The overall supervised loss function is a linear combination of *L_MinCut_* and *L_CE_* with a weight parameter *β*. To alleviate the instability issue of graph partitioning based on GNN, the soft TCN assignment module can be run multiple times to generate multiple soft TCN assignment matrices. For each of them, the hard assignment is conducted by assigning the cell (row) to the TCN (column) with the highest probability. Finally, an ensemble procedure is performed on those hard TCN assignments using the majority-voting strategy to determine the final TCN partition of the single-cell spatial map. **(b)** For an unsupervised learning task, CytoCommunity identifies TCNs for each tissue image individually. **(c)** For a supervised learning task, using a dataset of tissue images associated with different conditions as the input, CytoCommunity enables *de novo* identification of condition-specific TCNs under the supervision of image labels (e.g. TCNs colored by dark cyan or spring green).

### Performance evaluation using single-cell spatial proteomics data

To evaluate the performance of CytoCommunity, we applied it to a spatial proteomics dataset of mouse spleen generated using the Co-Detection by Indexing (CODEX) technology ^14^ and compared with four state-of-the-art methods including Spatial-LDA ^9^, STAGATE ^8^, BayesSpace ^5^ and stLearn ^6^ (Online Methods). This dataset includes CODEX images of three healthy mouse spleen samples (named as BALBc-1, BALBc-2 and BALBc-3). On average, each image contains 81,760 cells covering 27 cell types (Fig. 2a). The images are manually annotated by the authors with four known tissue compartments of the spleen: red pulp, marginal zone, B-cell zone and periarteriolar lymphoid sheath (PALS) (Fig. 2b). Here we regard these tissue compartments as ground-truth TCNs. We evaluate the agreement of predicted TCNs and the ground truth using three performance metrics: accuracy, normalized mutual information (NMI) and adjusted Rand index (ARI) (Online Methods). Overall, all five methods can identify the PALS compartment accurately. However, compared to CytoCommunity, the other four methods performed poorly in distinguishing marginal zone, red pulp and B-cell zone, especially marginal zone which was hardly identified by the other four methods (Fig. 2c). Quantitatively, CytoCommunity also achieved the highest accuracy, NMI and ARI values across the three images (Fig. 2d). In conclusion, CytoCommunity has substantially improved performance over representative state-of-the-art methods when comparing identified TCNs with manually annotated tissue compartments.

**Figure 2.**
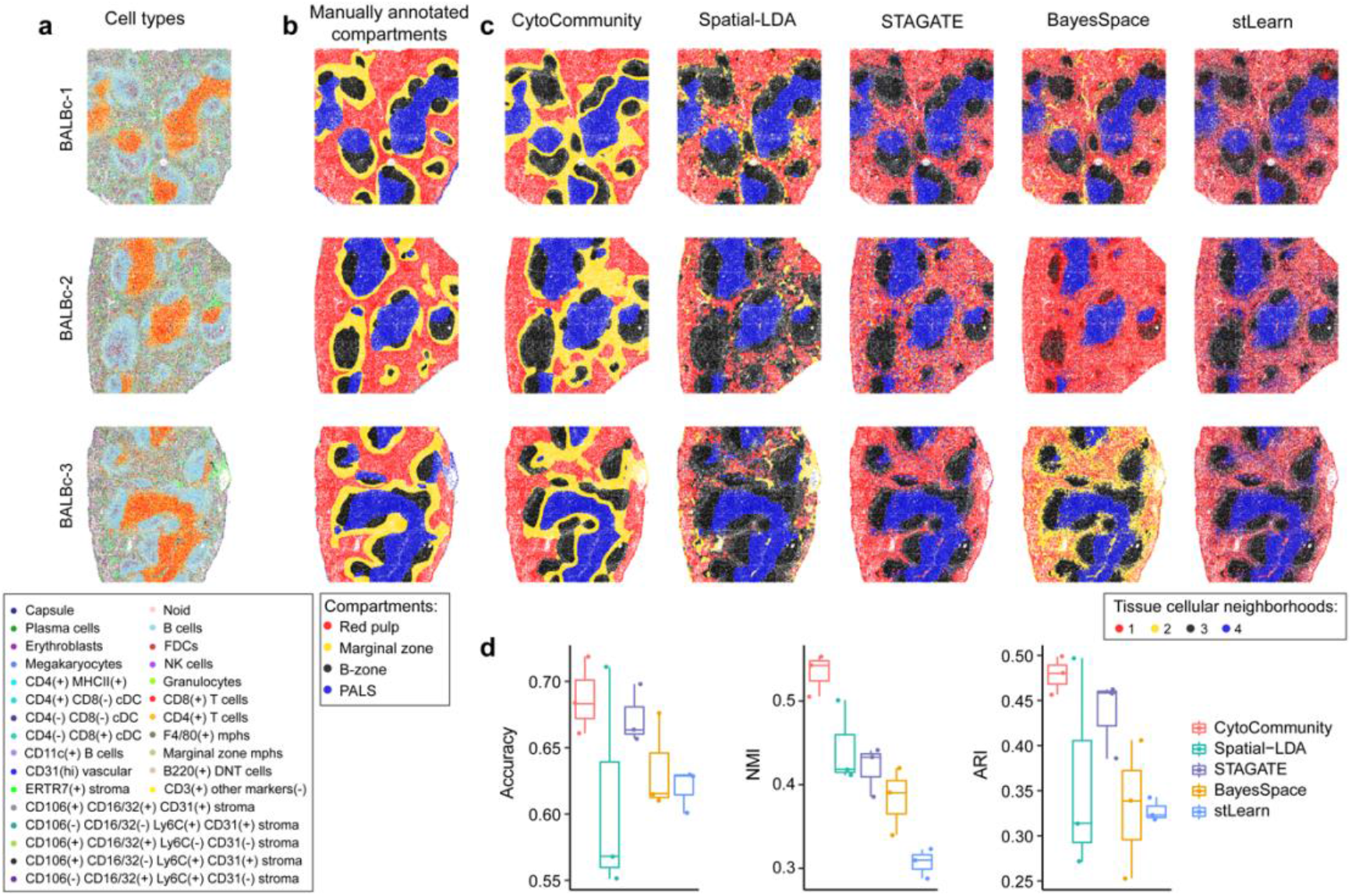
Performance evaluation of the CytoCommunity algorithm using single-cell spatial proteomics data. **(a, b)** Three single-cell spatial maps, BALBc-1, BALBc-2 and BALBc-3, generated from healthy mouse spleen samples by the CODEX technology. Cells are colored based on cell type annotation (a) or manual tissue compartment annotation (b) from the original study ^14^. **(c)** Tissue cellular neighborhoods (TCNs) identified by CytoCommunity, Spatial-LDA, STAGATE, BayesSpace and stLearn. **(d)** Boxplots of accuracy, normalized mutual information (NMI) and adjusted Rand index (ARI) values computed by comparing detected TCNs with manually annotated tissue compartments. Each dot in the boxplot represents the performance on a given single-cell spatial map.

### Performance evaluation using single-cell spatial transcriptomics data

The evaluation above focused on identifying large tissue compartments. To further evaluate the performance of CytoCommunity on detection of TCNs of smaller sizes, we applied it to a spatial transcriptomic dataset of healthy mouse hypothalamic preoptic region generated using the Multiplexed Error-Robust Fluorescence in situ Hybridization (MERFISH) technology ^15^. This dataset includes five MERFISH images with 18 manually annotated hypothalamic nuclei regions (Fig. 3a). In neuroanatomy, a nucleus is a group of neurons having similar connections and functions. Hence, we treated these manually annotated nuclei as gold-standard TCNs in the performance evaluation. On average, each image contains 5,352 cells that were assigned to nine cell types by the authors ^15^ (Fig. 3b). As shown in Fig. 3a, a prominent tissue architectural feature of the preoptic region is the symmetry of various types of nuclei. We found that CytoCommunity can identify multiple symmetric and coherent TCNs that agree with the manually outlined nuclei (Fig. 3c). For example, symmetric BNST (bed nucleus of the stria terminalis), MPA (medial preoptic area) and MPN (medial preoptic nucleus) regions were identified in all five images. We also identified symmetric VLPO (ventrolateral preoptic nucleus) regions in “Bregma-0.04”, “Bregma+0.06” and “Bregma+0.16” images, symmetric SHy (septohypothalamic nucleus), AVPe (anteroventral periventricular nucleus) and VMPO (ventromedial preoptic nucleus) regions in image “Bregma+0.06” and symmetric PaAP (paraventricular hypothalamic nucleus) regions in image “Bregma+0.26”. Besides these symmetric domains, central ACA (anterior commissure), Pe (periventricular hypothalamic nucleus) and MnPO (median preoptic nucleus) domains were also identified. In contrast, a number of manually annotated nuclei cannot be identified by the other four methods (unlabeled TCNs in the figure legend). Among those that can be identified, many are intermixed without clear boundary between them and lacks clear symmetry (Fig. 3c). In conclusion, CytoCommunity has substantially improved performance over state-of-the-art methods when comparing identified TCNs with manually annotated, complex tissue functional regions of variable sizes.

**Figure 3.**
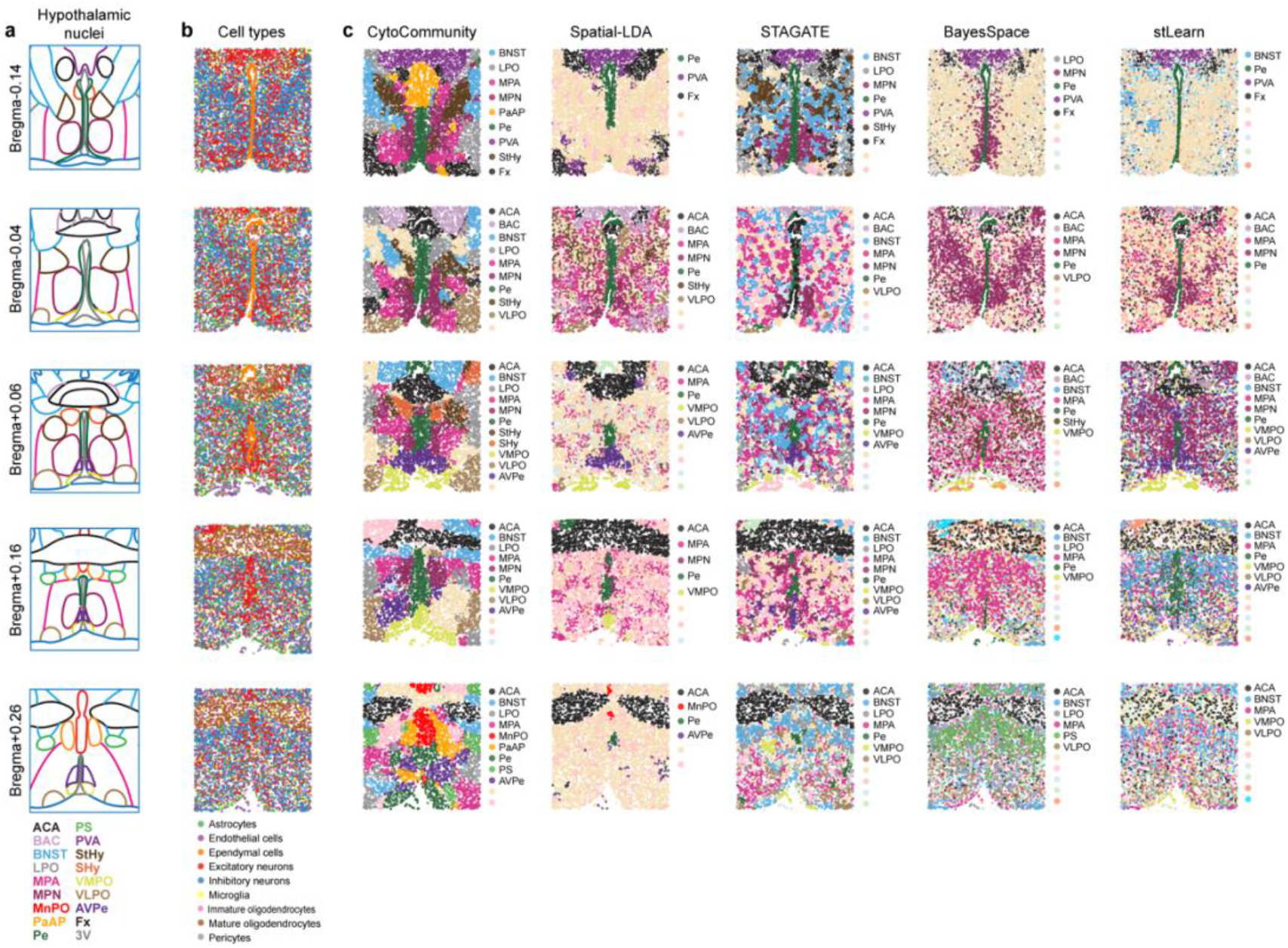
Performance evaluation of the CytoCommunity algorithm using single-cell spatial transcriptomics data. **(a)** Five single-cell spatial images, Bregma-0.14, Bregma-0.04, Bregma+0.06, Bregma+0.16 and Bregma+0.26, of the mouse hypothalamic preoptic region generated by the MERFISH technology. Bregma distance is given for each imaged brain section. 9, 10, 12, 12 and 11 hypothalamic nuclei/regions in the images are manually outlined by the authors ^15^. **(b)** Cells in the five images are colored based on the cell type annotation from the original study ^15^. **(c)** Tissue cellular neighborhoods (TCNs) identified by CytoCommunity, Spatial-LDA, STAGATE, BayesSpace and stLearn are labeled and colored based on the most similar manually annotated nuclei regions. TCNs without labels cannot be matched to the manual annotation. ACA, anterior commissure; BAC, bed nucleus of the anterior commissure; BNST, bed nucleus of the stria terminalis; LPO, lateral preoptic area; MPA, medial preoptic area; MPN, medial preoptic nucleus; MnPO, median preoptic nucleus; PaAP, paraventricular hypothalamic nucleus; Pe, periventricular hypothalamic nucleus; PS, parastrial nucleus; PVA, paraventricular thalamic nucleus; StHy, striohypothalamic nucleus; SHy, septohypothalamic nucleus; VMPO, ventromedial preoptic nucleus; VLPO, ventrolateral preoptic nucleus; AVPe, anteroventral periventricular nucleus; Fx, fornix; 3V, third ventricle.

### Altered tumor-immune interactions within and between tissue cellular neighborhoods in low- versus high-risk colorectal cancer patients

To demonstrate the utility of CytoCommunity for *de novo* identification of condition-specific TCNs by supervised learning, we applied it to a CODEX dataset generated using samples from 17 low-risk (characterized by “Crohn’s-like reaction”, CLR) and 18 high-risk (characterized by diffuse inflammatory infiltration, DII) colorectal cancer patients ^3^. The CLR patient group was reported to have significantly better overall survival than the DII patient group (log-rank test p = 0.002) ^3^. The dataset consists of 68 and 72 CODEX images from the CLR and DII patients, respectively (four images per patient). Using 10 sets of 10-fold cross-validation, we found that CytoCommunity nearly perfectly classified the images into the two patient groups with an average area under the receiver operating characteristic curve of 0.99 (Fig. 4a). We next investigated the 10 TCNs identified in the 140 CODEX images using supervised learning and found that the cell type enrichment scores (Online Methods) in those TCNs are significantly correlated (Pearson correlation coefficient (PCC) = 0.69, Fig. 4b and 4c), which indicates that the identified TCNs of the two patient groups have similar cell type composition. However, we also found multiple cell types that are enriched in CLR or DII-specific TCNs. For example, B cells are significantly enriched in TCN-5 in CLR patients but not enriched in any TCN in DII patients (Fig. 4b; Supplementary Fig. 1a), which is consistent with the presence of B cell-enriched tertiary lymphoid structures (TLSs) in CLR patient samples but absence of TLSs in DII patient samples ^3^. On the contrary, granulocytes are significantly enriched in TCN-7 in DII patients but not enriched in TCNs in CLR patients (Fig. 4b; Supplementary Fig. 1b), which is consistent with the previously reported critical role of granulocytes in DII patients ^3^. Interestingly, tumor cells and vascular smooth muscle cells are enriched in more TCNs in the DII group than in the CLR group (Fig. 4b), suggesting that the two cell types are spatially abundant in high-risk cancer patients.

**Figure 4.**
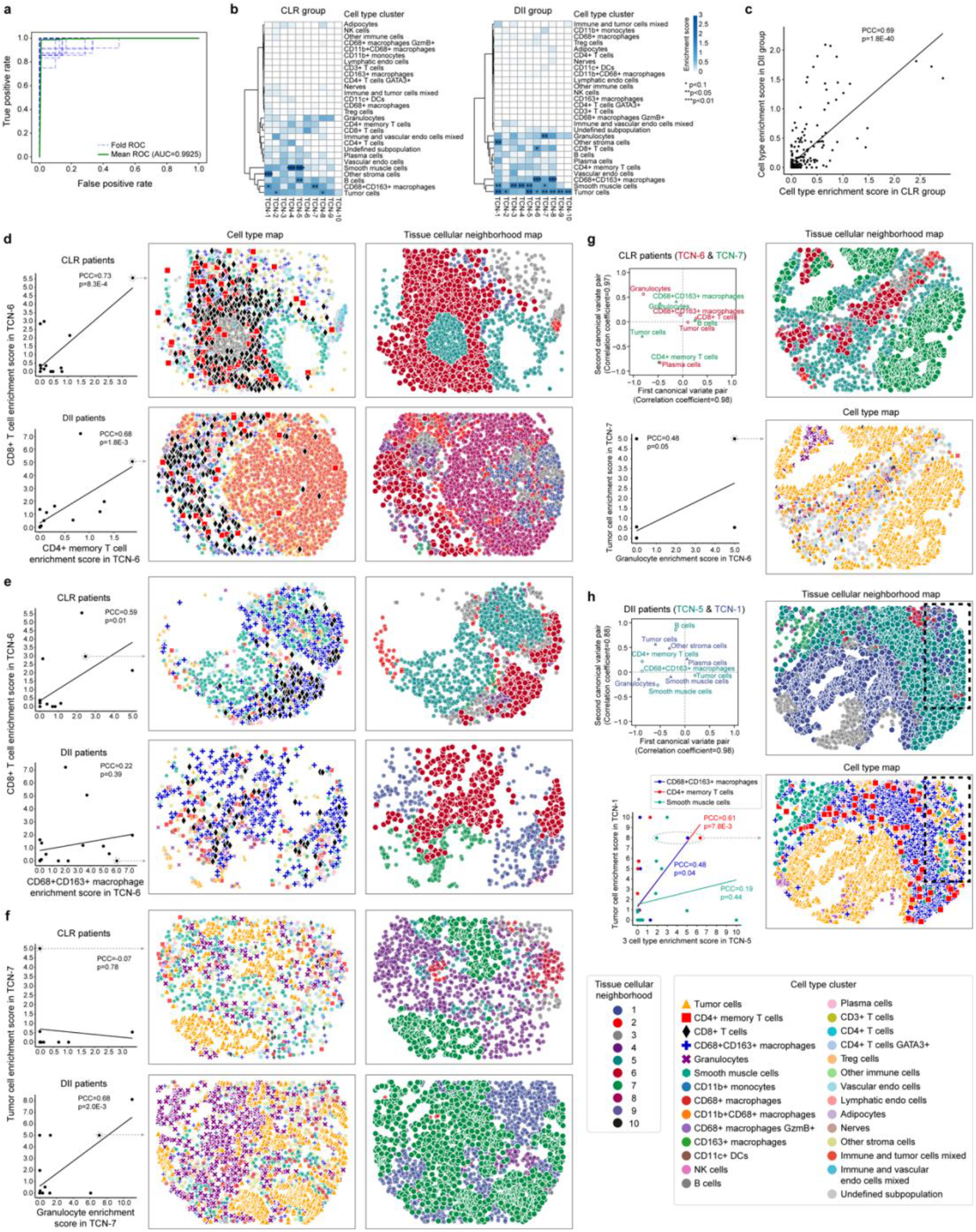
Coordinated tumor and immune cell type distributions within and between tissue cellular neighborhoods in colorectal cancer. **(a)** Receiver operating characteristic (ROC) curves for image label prediction based on 10 sets of 10-fold cross-validations. Blue dashed line, ROC curve for each fold of cross-validation. Green solid line, ROC curve based on the mean values of the 10 sets of 10-fold cross-validations. **(b)** Heatmaps of average enrichment scores of each cell type in each identified tissue cellular neighborhood (TCN) across all images of CLR and DII patient samples. Cell type enrichment score is defined as −log_10_(*P*-value). *P*-values were computed using hypergeometric test and adjusted using Benjamini-Hochberg method ^43^. **(c)** Correlation of average cell type enrichment scores in all identified TCNs between CLR and DII patients. **(d-f)** Correlation of the enrichment scores of two indicated cell types in TCN-6 or TCN-7 in each patient group (left panels). Representative cell type and TCN maps (middle and right panels) are based on patient samples indicated by a dashed circle in the scatter plots. **(g, h)** Significant canonical correlation (permutation test p-value < 0.05) between two TCNs in the CLR **(g)** and DII **(h)** patient groups. Shown are scatter plots of normalized weights of five cell types (observed variable) in each TCN in the first two canonical variate pairs (top left panels). Correlation of the enrichment scores of dominant cell types in the first canonical variate pair (bottom left panels), and representative cell type and TCN maps (right panels) are also shown. Black dashed rectangles in the cell type and TCN maps in **(h)** are used to highlight the colocalization of smooth muscle cells in TCN-5 and tumor cells in TCN-1. For all scatter plots, regression lines, Pearson correlation coefficients (PCC) and corresponding p-values are shown. For clarity, cells of studied types and TCNs are shown in large size without transparency in all cell type and TCN maps.

Besides the enrichment of individual cell types in TCNs, we also investigated the coordination of cell types within and between TCNs to better understand cell-cell communication in the tissue microenvironment. As an example of within-TCN cell type communication shared by the two patient groups (Supplementary Fig. 2), the enrichment of CD4+ memory T cells (red square) is significantly correlated with CD8+ T cell (black diamond) enrichment in TCN-6 in both CLR (PCC = 0.73) and DII (PCC = 0.68) patients (Fig. 4d, left panel). Consistent with the cell type and TCN maps from the patients with high enrichment scores, we observed that the two cell types are intermixed with each other in TCN-6 (Fig. 4d, middle and right panels). We also found CLR-specific (Supplementary Fig. 2a) and DII-specific (Supplementary Fig. 2b) cell type associations within TCNs. For instance, the enrichment of CD68+CD163+ macrophages (blue plus) is significantly correlated with CD8+ T cell enrichment in TCN-6 in CLR patients (PCC = 0.59) but not in DII patients (Fig. 4e), suggesting that double positive macrophages might have anti-tumor effects by promoting CD8+ T cell infiltration to improve survival of CLR patients. Similar cell type communication between the double positive macrophage and CD8+ T cell was recently reported in human lung cancer ^16^. As an opposite example, the enrichment of granulocytes (purple cross) and tumor cells in TCN-7 has significantly correlation (PCC = 0.68) in DII patients but not in CLR patients (Fig. 4f, left panel). Consistent with the corresponding cell type and TCN maps, we observed that a large number of granulocytes either intermix with or are spatially close to tumor cells in the DII group while much smaller number of granulocytes are near tumor cells in the CLR group (Fig. 4f, middle and right panels). This is in line with previous studies that neutrophils have tumor-promoting effects to impact cancer patient survival ^17, 18^.

To investigate communication between different TCNs, we conducted canonical correlation analysis of TCN pairs (Online Methods). We found substantial differences in significant canonical correlations of TCNs (permutation test p < 0.05) between CLR (Supplementary Fig. 3a) and DII patients (Supplementary Fig. 3b). As an example of significant between-TCN associations that are specific to CLR patients, granulocytes in TCN-6 and tumor cells in TCN-7 are two dominant cell types (observed variables) in the first canonical variate pair (Fig. 4g, top left panel). Without consideration of other cell types, granulocytes and tumor cells in the two TCNs have a statistically significant correlation (PCC = 0.48, p = 0.05) (Fig. 4g, bottom left panel), suggesting a potential interaction between this two cell types across TCNs. Consistent with the corresponding cell type and TCN maps, we observed that a small number of granulocytes enriched in TCN-6 are close to tumor cells enriched in TCN-7 (Fig. 4g, right panel). Such between-TCN communication in CLR patients is presumably different from the within-TCN communication between the same two cell types in DII patients (Fig. 4f), again supporting the pro-tumor role of granulocytes in DII patients.

Another interesting example of between-TCN communication regarding the DII group is the significant association between TCN-5 and TCN-1, in which CD68+CD163+ macrophages, CD4+ memory T cells, vascular smooth muscle cells, granulocytes and tumor cells are dominant cell types in the first canonical variate pair (Fig. 4h, top left panel). By examining the pair-wise correlation of these cell types, we found that double positive macrophages and CD4+ memory T cells in TCN-5 are significantly correlated with tumor cells in TCN-1 (PCC = 0.48 and 0.61, respectively; Fig. 4h, bottom left panel). Although vascular smooth muscle cells do not have significant correlation with tumor cells, the two cell types are co-enriched in the two TCNs in multiple DII patients (Fig. 4h, bottom left panel). From the corresponding cell type and TCN maps, we observed that the double positive macrophage/CD4+ memory T cell/vascular smooth muscle cell-enriched TCN-5 is spatially adjacent to the tumor cell-enriched TCN-1 (Fig. 4h, right panel), suggesting an unexpected cancer-promoting effects of these three cell types. As supporting evidence, extensive studies have demonstrated that tumor associated macrophages have functional plasticity that show both pro- and anti-tumor activities dependent on their microenvironment ^19, 20^. CD4+ memory T cells produce interleukin-22, which is induced by cancer cells to promote tumor growth in breast and lung cancers ^21^. Previous studies also reported the critical role of vascular smooth muscle cells in tumor angiogenesis and metastasis ^22^, which is consistent with our observation that a small group of tumor cells in TCN-1 reside with vascular smooth muscle cells in TCN-5 (Fig. 4h, right panel).

### Altered tumor-stromal interactions within and between tissue cellular neighborhoods in low- versus high-risk breast cancer patients

To further evaluate the ability of CytoCommunity to discover condition-specific TCNs using different data modalities, we applied it to another spatial proteomics dataset of breast cancer generated using the imaging mass cytometry technology ^10^. Based on the median overall survival, we stratified the 79 breast cancer patients into low- and high-risk groups with significant survival difference (log-rank test p < 0.0001; Fig. 5a). We identified nine TCNs in both low- and high-risk groups. By comparing their cell type enrichment scores (Fig. 5b and 5c), we found that TCNs in both groups have similar overall cell type composition (Fig. 5c) and are enriched for several types of fibroblasts (Fig. 5b), suggesting a critical role of fibroblasts in breast cancer prognosis. Specifically, we found that SMA^hi^ Vimentin^hi^ fibroblasts, also known as cancer associated fibroblasts (CAFs) that highly express alpha smooth muscle actin (SMA) and vimentin, are more enriched in TCNs of the high-risk group than those of the low-risk group (Fig. 5b; Supplementary Fig. 4a). Besides stromal cell types, we also found low- and high-risk-group specific TCNs, characterized by significant enrichment (p < 0.05) of CK^+^ HR^hi^ tumor cells (cells with positive expression of cytokeratins (CK) and high expression of hormone receptors (HR)) and CK^low^ HR^low^ tumor cells, respectively (Fig. 5b; Supplementary Fig. 4b and 4c). This is consistent with the previous report that these two tumor cell phenotypes are associated with good and poor prognosis, respectively ^10^.

**Figure 5.**
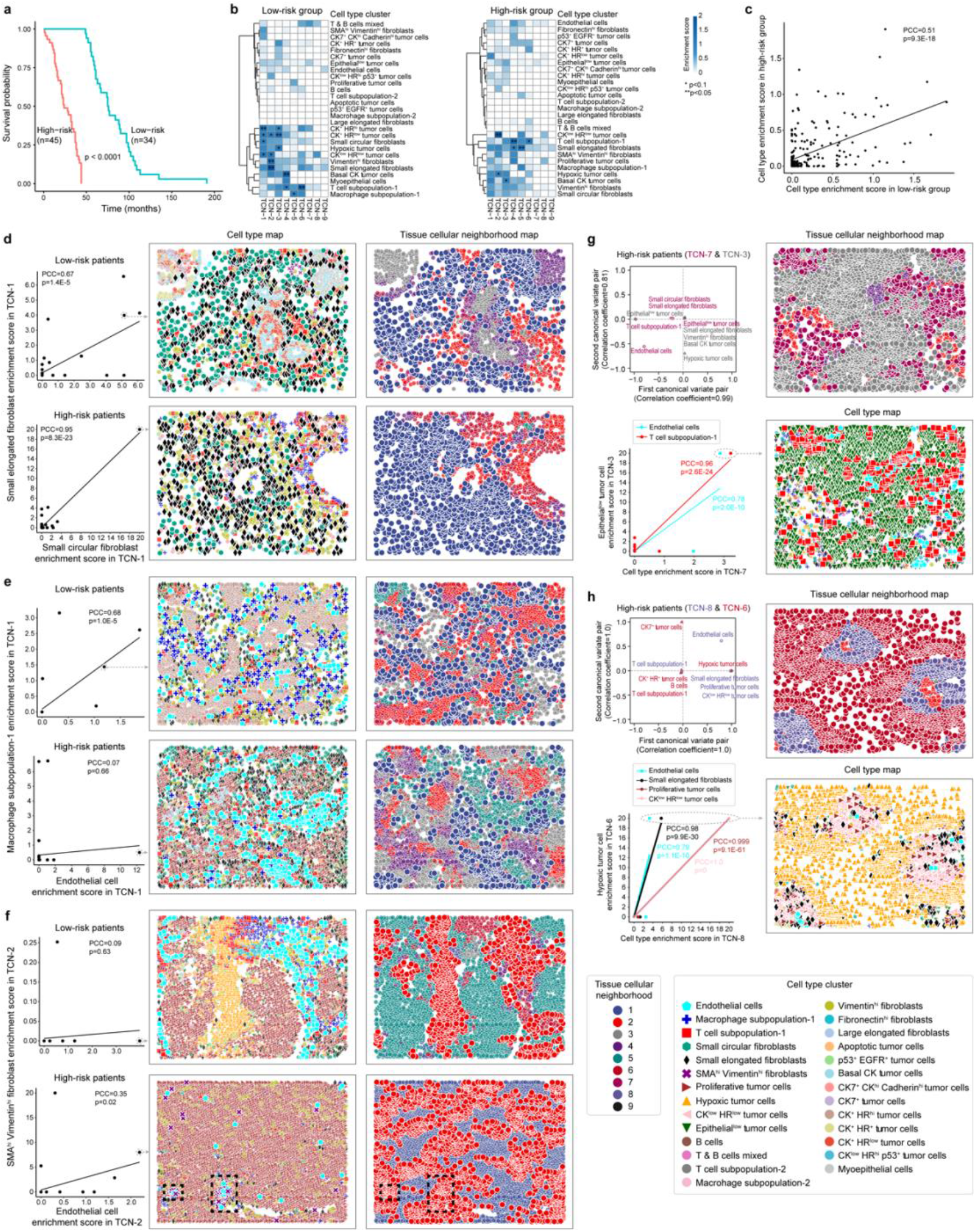
Coordinated tumor and stromal cell type distributions within or between tissue cellular neighborhoods in breast cancer. **(a)** Kaplan-Meier survival curves of 79 breast cancer patients who were classified into low-risk and high-risk groups based on their median overall survival time. *P*-value was computed using the log-rank test. **(b)** Heatmaps of average enrichment scores of each cell type in each identified tissue cellular neighborhood (TCN) across all images of low-risk and high-risk patient samples. Cell type enrichment score is defined as −log_10_(*P*-value). *P*-values were computed using hypergeometric test and adjusted using Benjamini-Hochberg method ^43^. **(c)** Correlation of average cell type enrichment scores in all identified TCNs between low-risk and high-risk patients. **(d-f)** Correlation of the enrichment scores of two indicated cell types in TCN-1 or TCN-2 in each patient group (left panels). Representative cell type and TCN maps (middle and right panels) are based on patient samples indicated by a dashed circle in the scatter plots. Black dashed rectangles in the cell type and TCN maps in **(f)** are used to highlight the colocalization of endothelial cells and SMA^hi^ vimentin^hi^ fibroblasts in TCN-2. **(g, h)** Significant canonical correlation (permutation test p-value < 0.1) between TCN-7 and TCN-3 **(g)** and between TCN-8 and TCN-6 **(h)** in the high-risk patient groups. Shown are scatter plots of normalized weights of five cell types (observed variable) in each TCN in the first two canonical variate pairs (top left panels). Correlation of the enrichment scores of dominant cell types in the first canonical variate pair (bottom left panels), and representative cell type and TCN maps (right panels) are also shown. For all scatter plots, regression lines, Pearson correlation coefficients (PCC) and corresponding p-values are shown. For clarity, cells of studied types and TCNs are shown in large size without transparency in all cell type and TCN maps.

Regarding cell type associations within TCNs (Supplementary Fig. 5), we found that two normal fibroblast types, small circular (green hexagon) and elongated (black diamond) fibroblasts, are significantly correlated in TCN-1 in both low- (PCC = 0.67) and high-risk (PCC = 0.95) patients (Fig. 5d, left panel). Correspondingly, we observed that these two fibroblast types are intermixed in TCN-1 in patients with high enrichment scores (Fig. 5d, middle and right panels). As examples of low- (Supplementary Fig. 5a) and high-risk-specific (Supplementary Fig. 5b) within-TCN cell-cell communications, we found that endothelial cell (cyan pentagon) enrichment is significantly correlated with macrophage (blue plus) enrichment in TCN-1 in low-risk patients (PCC = 0.68; Fig. 5e, left panel) but correlated with CAF (purple cross) enrichment in TCN-2 in high-risk patients (PCC = 0.35; Fig. 5f, left panel). This observation implies that differential interactions between endothelial cells, macrophages and fibroblasts can lead to different patient outcomes ^23, 24^. From representative cell type and TCN maps, we observed that endothelial cells are intermixed with macrophages in TCN-1 in the low-risk patient only (Fig. 5e, middle and right panels) but with CAFs in TCN-2 in the high-risk patient only (Fig. 5f, middle and right panels).

Next, we investigated between-TCN communications by canonical correlation analysis. Interestingly, we only found significant TCN associations involving tumor-stromal interactions in high-risk patients (Supplementary Fig. 6). For example, we observed that endothelial cell (cyan pentagon) and T cell (red square)-dominated TCN-7 is associated with TCN-3 dominated by epithelial^low^ tumor cells (green upside-down triangle) that are undergoing epithelial-mesenchymal-transition (EMT) (Fig. 5g). As supporting evidence, previous studies showed that both endothelial cells and T cells can induce EMT of cancer cells, resulting in poor clinical outcomes ^25–29^. Another interesting example is the significant correlation between hypoxic tumor cell (orange triangle)-dominated TCN-6 and TCN-8 that mainly consists of endothelial cells (cyan pentagon), small elongated fibroblasts (black diamond), proliferative (brown right-pointing triangle) and CK^low^ HR^low^ (pink left-pointing triangle) tumor cells (Fig. 5h). Proliferative tumor cells in TCN-8 probably lead to reduction in the oxygen supply, promoting the formation of a hypoxic tumor microenvironment ^30^. Hypoxia enables the expansion of aggressive tumor clones ^30^ (represented by the cohesive hypoxic tumor mass in TCN-6) and can also inhibit hormone receptor expression ^31^, probably contributing to the CK^low^ HR^low^ tumor phenotype in TCN-8. Hypoxic microenvironment can also support the transformation of tissue-resident fibroblasts to CAFs and endothelial cell-mediated neovascularization ^30^ in TCN-8. All these hypoxia-induced events are associated with unfavorable prognosis in breast cancer.

## Discussion

We introduce a deep graph learning approach, CytoCommunity, for identifying tissue cellular neighborhoods based on cell phenotypes and cell spatial distributions. CytoCommunity formulates the TCN identification as a community detection problem on node-attributed cell-cell spatial proximity graphs. Since most traditional community detection algorithms focus only on graph topology to find densely-connected subgraphs and cannot explicitly deal with node attributes ^32^, CytoCommunity employs a minimum cut-based GNN model to learn optimal TCN assignments of cells (nodes) from cell type information (node attributes). Like previous methods ^2, 4–9^, CytoCommunity can be applied in an unsupervised fashion to identify TCNs in individual images. More importantly, it is the first TCN-detection method that can be used in a supervised fashion for *de novo* identification of condition-specific TCNs and prediction of tissue image labels and therefore facilitating the discovery of more physiologically or clinically relevant tissue structures. This unique characteristics of CytoCommunity is attributed to the usage of graph pooling that preserves TCN partition information in the embedding representation of the whole image and thus addresses the TCN alignment across images by training an end-to-end model for image classification. It is worth noting that TCN identification under the supervision of image labels can be considered as a weakly supervised graph partitioning problem, representing an interesting research topic in graph learning.

Successful identification of large (splenic compartments) and small (hypothalamic nuclei) TCNs by unsupervised CytoCommunity suggest that information about cell types and their spatial distributions are sufficient to determine the functional units in tissues without using gene or protein expression features. By applying supervised CytoCommunity to risk-stratified cancer tissue images, we identified both shared and specific TCNs, such as a low-risk-specific TCN enriched for B cells, corresponding to the well-known tertiary lymphoid structure (TLS) associated with favorable prognosis. We also found granulocyte- and cancer-associated fibroblast-enriched TCNs in high-risk colorectal and breast cancer patients, respectively. By comparative analysis of TCNs, we revealed multiple altered tumor-immune and tumor-stromal coordination patterns within and between TCNs in low-versus high-risk cancer patients.

We believe that the success of CytoCommunity can be attributed to two main features. First, it leverages a GNN model with a theoretically grounded minimum cut-based loss function ^13^ for soft TCN assignment learning, generating more accurate and stable graph partitioning results than other pooling-capable GNN models such as DiffPool ^33^, which employs heuristic loss functions to learn the soft assignments. Second, CytoCommunity uses cell types as initial cell features, probably leading to a better measurement of functional similarity between cells than using noisy gene or protein expression data directly. Cell type identification is typically the first crucial task in single-cell data analysis and often needs sophisticated tools ^34–36^ as well as expert knowledge. Therefore, cell type annotation should be directly utilized in a specialized TCN detection method rather than starting with expression data. CytoCommunity encodes cell types in categorical vector space and thus has scalability to incorporate more heterogenous categorical data, such as cell states ^37^, into the initial cell feature vectors for inferring TCNs.

Due to the use of cell type information, the current version of CytoCommunity is not applicable to spatial transcriptomics data with spot resolution ^38,39^. To address this issue, cell type composition at each spot can be first estimated by deconvolution methods ^40, 41^. Then, a spot-spot proximity graph with inferred cell type fractions as node attributes can be constructed as the input to CytoCommunity.

In summary, with the rapid growth of single-cell spatial maps, CytoCommunity represents a powerful and scalable method for identifying condition-specific TCNs. TCNs directly learned from cell types can facilitate their function interpretation and discovery of cell-cell communications within the tissue microenvironment.

## Online Methods

### Unsupervised model for identification of tissue cellular neighborhoods

The CytoCommunity algorithm for identifying tissue cellular neighborhoods (TCNs) consists of two components: a soft TCN assignment learning module and a TCN ensemble procedure to determine the final robust TCNs (Fig. 1a). As the first component, given a single-cell spatial map with cell type annotation and cell location data, an undirected K-nearest neighbor (KNN) graph with node attribute (cell type) is constructed. In the graph, a cell is represented by a node and its cell type information (categorical data) is represented by a node attribute vector using one-hot encoding (Fig. 1a, top panel). Specifically, we first construct a directed KNN graph by connecting each node to its KNNs based on Euclidean distance calculated using cell spatial coordinates. Then, the underlying undirected graph without self-edges of the directed KNN graph was used as the input to the graph neural network (GNN). Since each spatial omics dataset is measured from the same tissue type and using the same technology, we set the value of K in the KNN graphs as the square root of the average number of cells across images in the dataset.

For the undirected KNN graph with *n* nodes, we employ a basic GNN model ^42^ with the ReLU activation function to generate a node embedding matrix *X ∈ ℝ^n×d^*, where each row is a learned *d*-dimensional representation vector of a node defined as below.

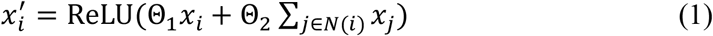

where 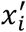 is an updated embedding vector of node *i*, which is calculated based on the previous representation of itself *x_i_* and its first-order neighborhood *N(i)* in the undirected KNN graph constructed above. *x_i_* is initialized with the node attribute vector. Θ_1_ and Θ_2_ are trainable parameter matrices in the GNN model. The value of *d* was empirically set to 128 for all datasets in this study.

Next, we use a fully-connected neural network with no hidden layers, also known as a linear layer, and the softmax activation function to transform the node embedding matrix *X ∈ ℝ^n×d^* to the soft TCN assignment matrix *S ∈ ℝ^n×c^*, which can be formulated as below.

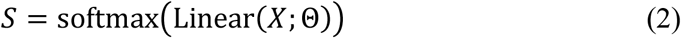

where each element in *S* represents the probability of a node (row) belonging to one of the *c* TCNs. *c* is a user-specified hyperparameter and represents the maximum number of TCNs to be detected. Next, we use the following graph minimum cut (MinCut)-based loss function ^13^ to optimize the matrix *S* in an unsupervised way.

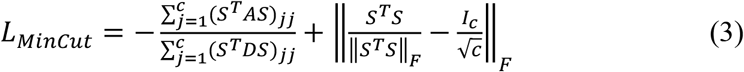

where *A ∈ {0,1}^n×n^* is the symmetric adjacency matrix derived from the undirected KNN graph. *D ∈ ℝ^n×n^* is a diagonal matrix (or the degree matrix), where each diagonal element is the sum of the corresponding row in *A*. The loss function *L_MinCut_* is the sum of two terms. The left term is used to address the normalized MinCut problem in graph theory with the objective of partitioning the graph into *c* disjoint connected components with similar sizes by removing the minimum number of edges. The right term encourages the soft TCN assignment matrix *S* to be orthogonal in order to make the TCN membership of each node unambiguous. ||·||*_F_* denotes the Frobenius norm. This loss function can be used alone for an unsupervised learning task, that is *L_Unsup_* = *L_MinCut_*, to identify TCNs for single-cell spatial omics images individually (Fig. 1b).

As the second component of the CytoCommunity algorithm, we attempt to obtain a robust graph partitioning result as the final TCNs by conducting a TCN ensemble procedure (Fig. 1a, bottom panel). Specifically, the soft TCN assignment learning module in the first component is run multiple times to generate multiple learned matrices ***S*.** For each of them, the hard assignment is performed by assigning the cell (row) to the TCN (column) with the highest probability. Then, we use the majority-voting strategy to conduct an ensemble procedure on those hard TCN assignments to determine the final TCN partition of the single-cell spatial map. To demonstrate the effectiveness of this ensemble-based approach, we performed a stability experiment on the challenging MERFISH dataset with multiple TCNs of small sizes to be detected (Supplementary Fig. 7). Specifically, we applied CytoCommunity with different number of runs of the first component to the MERFISH image for five times. We then conducted pair-wise comparisons among the TCN partitions generated by the five sets of runs. Our results showed that the TCN ensemble procedure based on 20 runs is sufficient to obtain a stable TCN partition.

### Supervised model for *de novo* identification of condition-specific tissue cellular neighborhoods

Given a dataset of multiple spatial omics images from different conditions, TCNs can be first identified for each image and then aligned across images for identifying condition-specific TCNs. However, TCN alignment is analogous to community alignment in graphs, which is NP-hard ^12^. To tackle this problem, we take advantage of graph pooling to generate an embedding representation of the whole graph that preserves the TCN partition information. Thus, by adapting the unsupervised graph partitioning model described above to a graph convolution and pooling-based graph classification framework, TCNs in different images are automatically aligned during soft TCN assignment learning, facilitating the identification of condition-specific TCNs (Fig. 1a, top panel). Specifically, after obtaining the soft TCN assignment matrix *S* by a graph convolution and a fully-connected layers, we additionally employ a graph pooling layer ^13, 33^ formulated as below to generate a coarsened adjacency matrix *A^Pooled^ ∈ ℝ^c×c^* and a matrix of embeddings *X^Pooled^ ∈ ℝ^c×d^* for the *c* pooled nodes in the coarsened graph. Note that this coarsened graph is a fully connected graph with each pooled node corresponding to a cluster of nodes in the original KNN graph and the weights of edges representing the connectivity strengths between clusters.

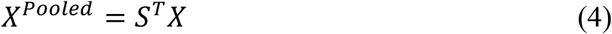

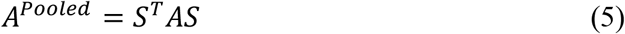

Then, we use *X^Pooled^* and *A^Pooled^* as inputs to another GNN same as described in Equation (1) to integrate pooled node features and their local neighborhood information in the coarsened graph, generating an updated embedding vector for each pooled node. The average across these new embedding vectors of the pooled nodes is considered as an embedding vector of the whole graph, which is in turn used as the input to a graph classifier implemented by two fully-connected layers with the softmax activation function. The overall supervised loss function is defined as follows.

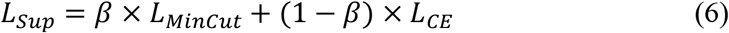

where *β* is a weight parameter to balance the minimum cut loss *L_MinCut_* used for graph partitioning as described above and the cross-entropy loss *L_CE_* used for graph classification. Trained with the joint loss function, this model is able to directly learn condition-specific TCNs under the supervision of graph (image) labels (Fig. 1c).

For both colorectal and breast cancer datasets in this study, we performed 10 sets of 10-fold cross-validation to evaluate prediction performance of the model and used 100 optimal soft TCN assignment matrices generated during cross-validation to conduct the TCN ensemble procedure for robust TCN identification. We empirically set *β* to 0.9 due to our emphasis on graph partitioning (i.e. TCN identification) and set the maximum number of TCNs that will be identified (i.e. the value of *c*) to 10.

### Running of published methods

We compared the performance of CytoCommunity with four other spatial domain detection methods, including Spatial-LDA ^9^, STAGATE ^8^, BayesSpace ^5^ and stLearn ^6^. As required by these methods, cell type annotation and cell spatial coordinates were used as inputs to Spatial-LDA, while protein or mRNA expression data and cell spatial coordinates were used as inputs to the other three methods. For benchmarking purpose, the number of TCNs to be detected were specified according to the manual annotation from the original studies ^14, 15^.

The Python package “spatial-lda (v0.1.3)” was applied to the CODEX dataset of mouse spleen and the MERFISH dataset of mouse hypothalamic preoptic region. By considering all cells as index cells, we first used *featurize_samples* and *make_merged_difference_matrices* functions for image featurization. Then, TCNs were detected using *spatial_lda.model.train* function with parameters max_dirichlet_iter=30 and max_dirichlet_ls_iter=30 for the CODEX dataset and default parameters for the MERFISH dataset. Note that the number of TCNs identified by this method may be fewer than the pre-specified number.

The Python package “STAGATE-pyG (v1.0.0)” was applied to both datasets. For each image, cell spatial neighbor network was constructed using *Cal_Spatial_Net* and *Stats_Spatial_Net* functions. The *train_STAGATE* function was then used to learn low-dimensional latent representations of cells, which were considered as inputs to the Louvain clustering algorithm for TCN detection. *scanpy.pp.neighbors* and *scanpy.tl.louvain* functions were used here with resolution=0.25 for all three CODEX images. In order to obtain the same number of TCNs as the manually outlined hypothalamic nuclei regions in each MERFISH image, the parameter resolution was set to 0.5, 0.45, 0.6, 0.62 and 0.76 for image Bregma-0.14, Bregma-0.04, Bregma+0.06, Bregma+0.16 and Bregma+0.26, respectively.

The R package “BayesSpace (v1.5.1)” was applied to the CODEX dataset with top 15 principal components (n.PCs) considered and all 30 protein markers as highly variable genes (n.HVGs) in the preprocessing function *spatialPreprocess*. TCNs were then identified using *spatialCluster* function with nrep=5000 and burn.in=100. For the MERFISH dataset, TCNs were identified using *spatialPreprocess* function with n.PCs=30 and n.HVGs=155 and *spatialCluster* function with nrep=10,000 and burn.in=100.

The Python package “stlearn (v0.4.0)” was applied to both datasets following the stLearn official tutorial “Working with MERFISH” https://stlearn.readthedocs.io/en/latest/tutorials/Read_MERFISH.html. The parameters were set to be n_comps=30, randome_state=0, and n_neighbors=50 in the preprocessing functions *stlearn.em.run_pca* and *stlearn.pp.neighbors* for both datasets. Then, TCNs were identified using *stlearn.tl.clustering.louvain* function with resolution=0.25 for the CODEX dataset. With respect to the MERFISH dataset, the parameter resolution was set to 0.35, 0.5, 0.8, 0.9 and 1.3 for image Bregma-0.14, Bregma-0.04, Bregma+0.06, Bregma+0.16 and Bregma+0.26, respectively.

### Quantitative performance evaluation using the CODEX dataset of mouse spleen

We used three metrics, accuracy, normalized mutual information (NMI) and adjusted Rand index (ARI) to quantitatively evaluate the performance of five compared methods. The ground-truth labels of cells among four known splenic compartments, red pulp, marginal zone, B-cell zone and periarteriolar lymphoid sheath (PALS), were obtained from the authors of the original study ^14^. For each CODEX image, NMI, ARI and accuracy were defined as below and computed using the R package “aricode (v1.0.0)”. Note that we assigned each identified TCN to the compartment with the largest number of cell matches.

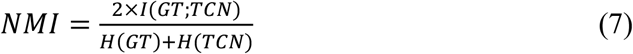

where *I (GT; TCN)* is the mutual information between ground-truth and predicted TCN labels of cells. *H(GT)* and *H(TCN)* are the entropies of ground-truth and predicted TCN labels, respectively.

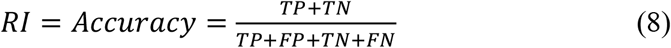

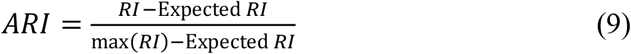

where the Rand index (RI) is computed based on true positives (TP), true negatives (TN), false positives (FP) and false negatives (FN) of the TCN predictions compared to the ground-truth labels of cells.

### Cell type enrichment score in tissue cellular neighborhoods

To quantitatively measure the composition of cell types in identified TCNs, we defined an enrichment score of each cell type in each TCN as −log_10_(*P*-value). The *P*-value was computed using hypergeometric test based on the following four numbers: (1) the number of cells of a given type in the TCN; (2) the total number of cells in the TCN; (3) the number of cells of the given type in the single-cell spatial omics image; (4) the total number of cells in the image. *P*-values were adjusted for multiple testing using the Benjamini-Hochberg method ^43^.

### Canonical correlation analysis of tissue cellular neighborhoods

To identify the associations among cell types located in different TCNs, we conducted canonical correlation analysis (CCA) of each TCN pair using cell type enrichment scores. Specifically, for each TCN, we selected five most variable cell types based on the standard deviation of enrichment scores across patient samples as observed variables of the TCN. Then, the canonical correlation model between each TCN pair was constructed using the *cc* function from the R package “CCA (v1.2.1)”. We also computed statistical significance p-values of canonical correlation coefficients using permutation test-based *p.perm* function from the R package “CCP (v1.2)”. To facilitate interpretation of CCA results, we further investigated correlations between dominant cell types identified based on their normalized weights in the first canonical variate pair to describe cell type communication patterns between TCNs.

### Cancer risk stratification based on survival analysis

For the spatial proteomics dataset of breast cancer generated using the imaging mass cytometry technology ^10^, we conducted a patient stratification into low- and high-risk groups based on the median overall survival of 79 deceased patients only. We did not consider censored patients since their overall survival time is unknown. The Kaplan-Meier survival curves and corresponding log-rank test p-value were computed using the R package “survival (v3.2-13)”.

## Supplementary Figure Legends

**Supplementary Figure 1.**
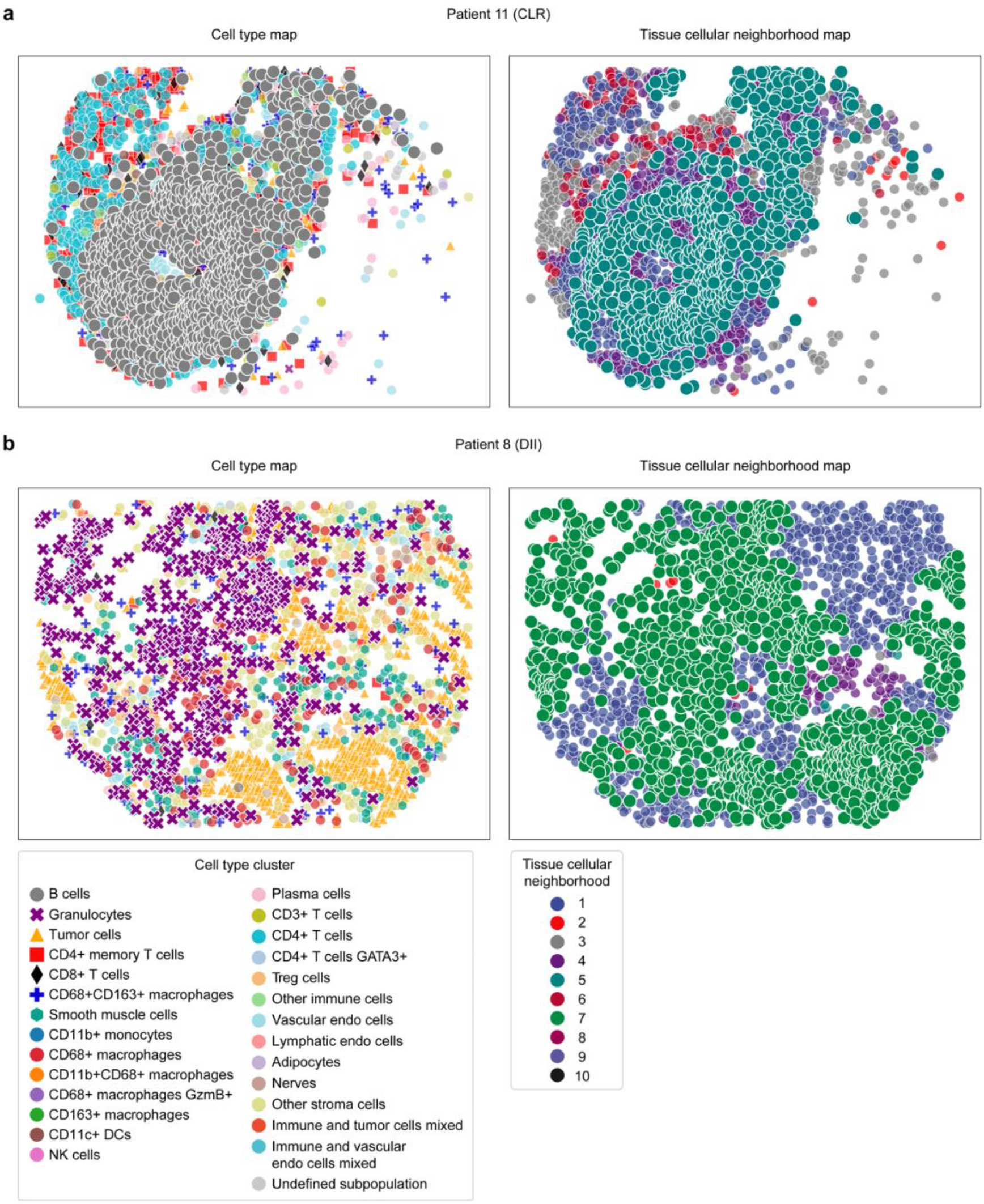
Representative tissue cellular neighborhoods specific to low- and high-risk colorectal cancer patients. **(a)** Cell type and tissue cellular neighborhood (TCN) maps of a representative CLR patient image with B cells enriched in TCN-5. **(b)** Cell type and TCN maps of a representative DII patient image with granulocytes enriched in TCN-7. For clarity, cells of studied types and TCNs are shown in large size without transparency in all cell type and TCN maps.

**Supplementary Figure 2.**
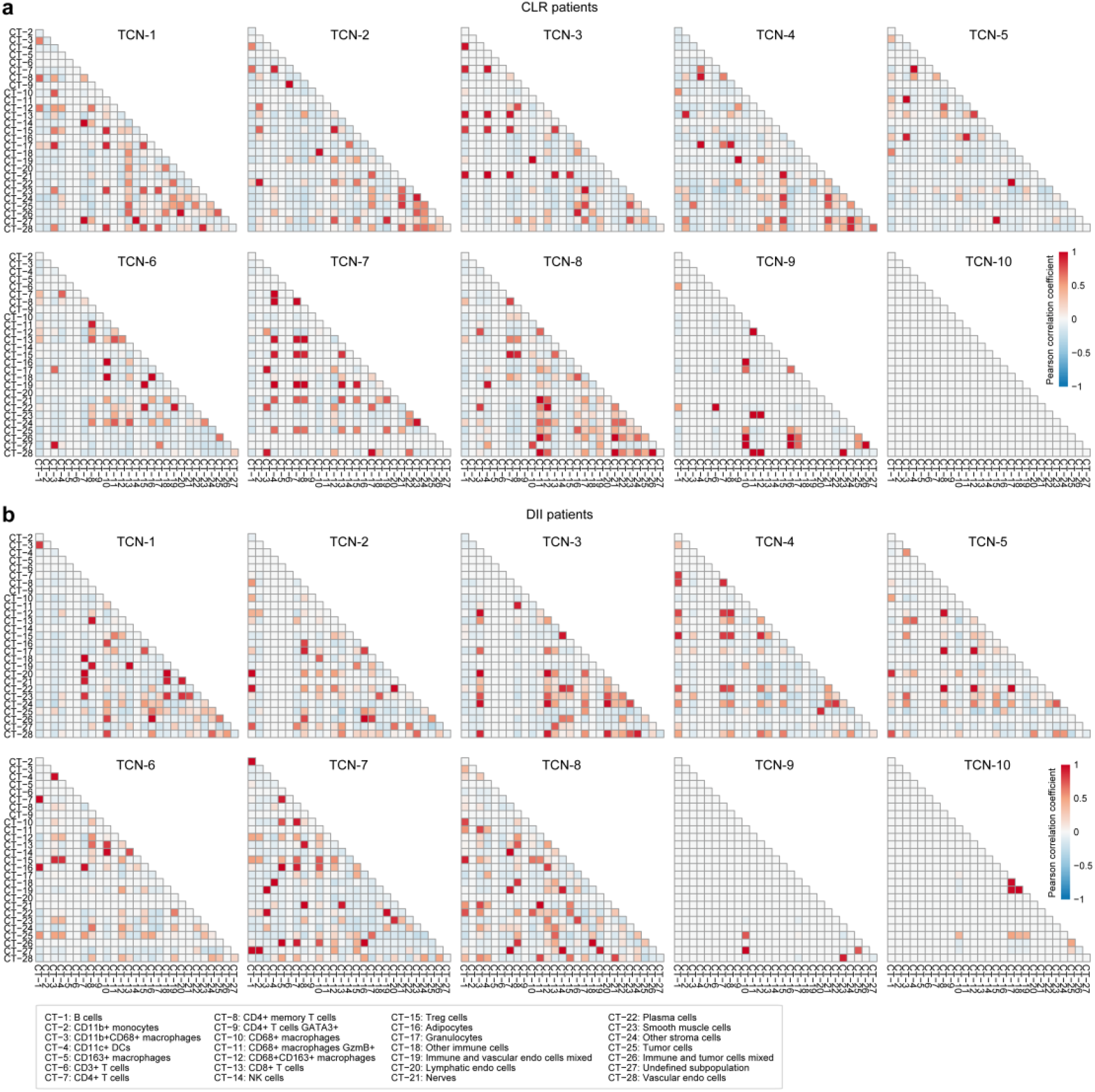
Cell type associations within tissue cellular neighborhoods in colorectal cancer. Heatmaps for correlations of the enrichment scores of any two cell types (CTs) within each of the 10 tissue cellular neighborhoods (TCNs) identified in CLR **(a)** and DII **(b)** patients.

**Supplementary Figure 3.**
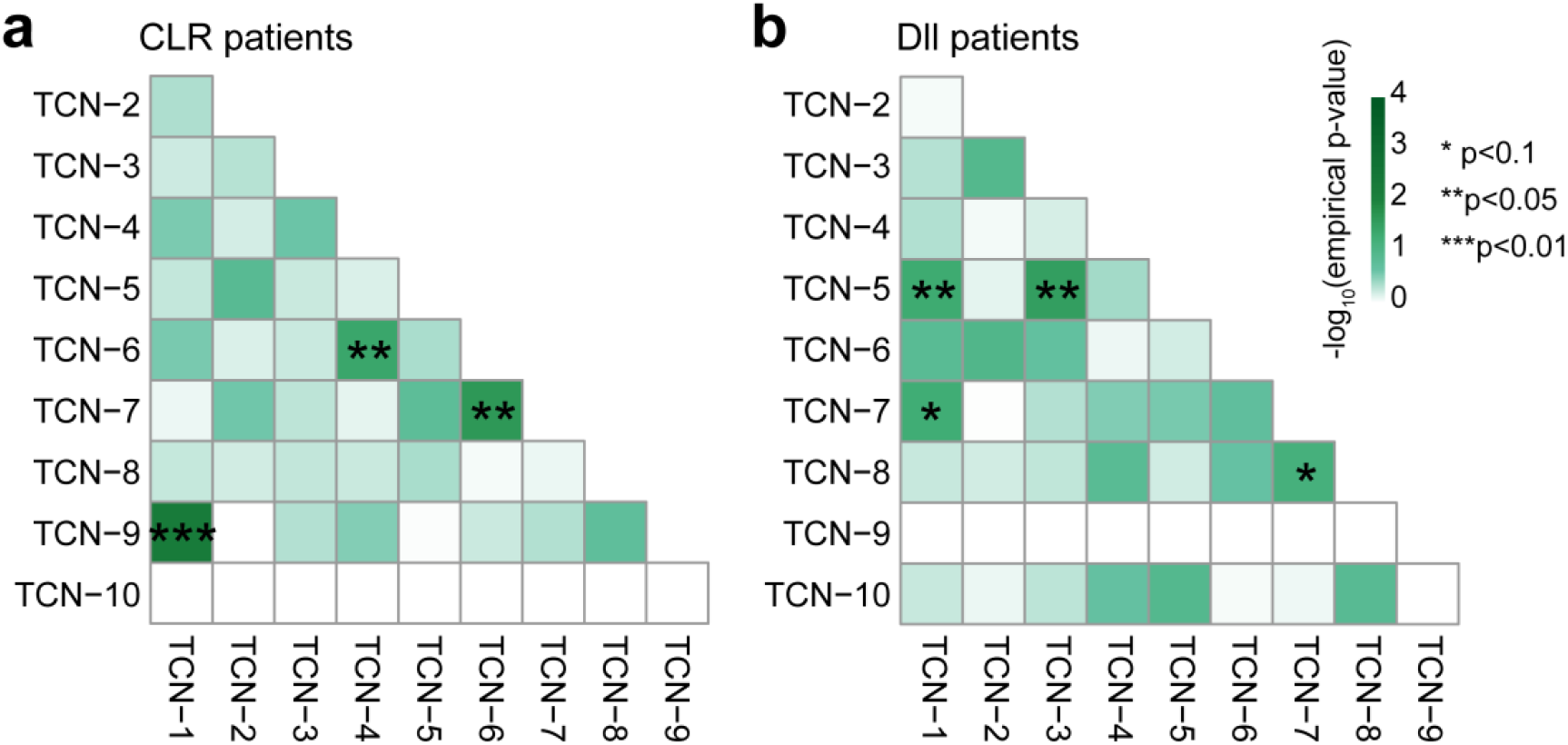
Cell type associations between tissue cellular neighborhoods in colorectal cancer. Heatmaps for empirical p-values of canonical correlation coefficients of the cell type enrichment scores of tissue cellular neighborhood (TCN) pairs in CLR **(a)** and DII **(b)** patient groups. P-values were computed using permutation test-based *p.perm* function from the R package “CCP (v1.2)”.

**Supplementary Figure 4.**
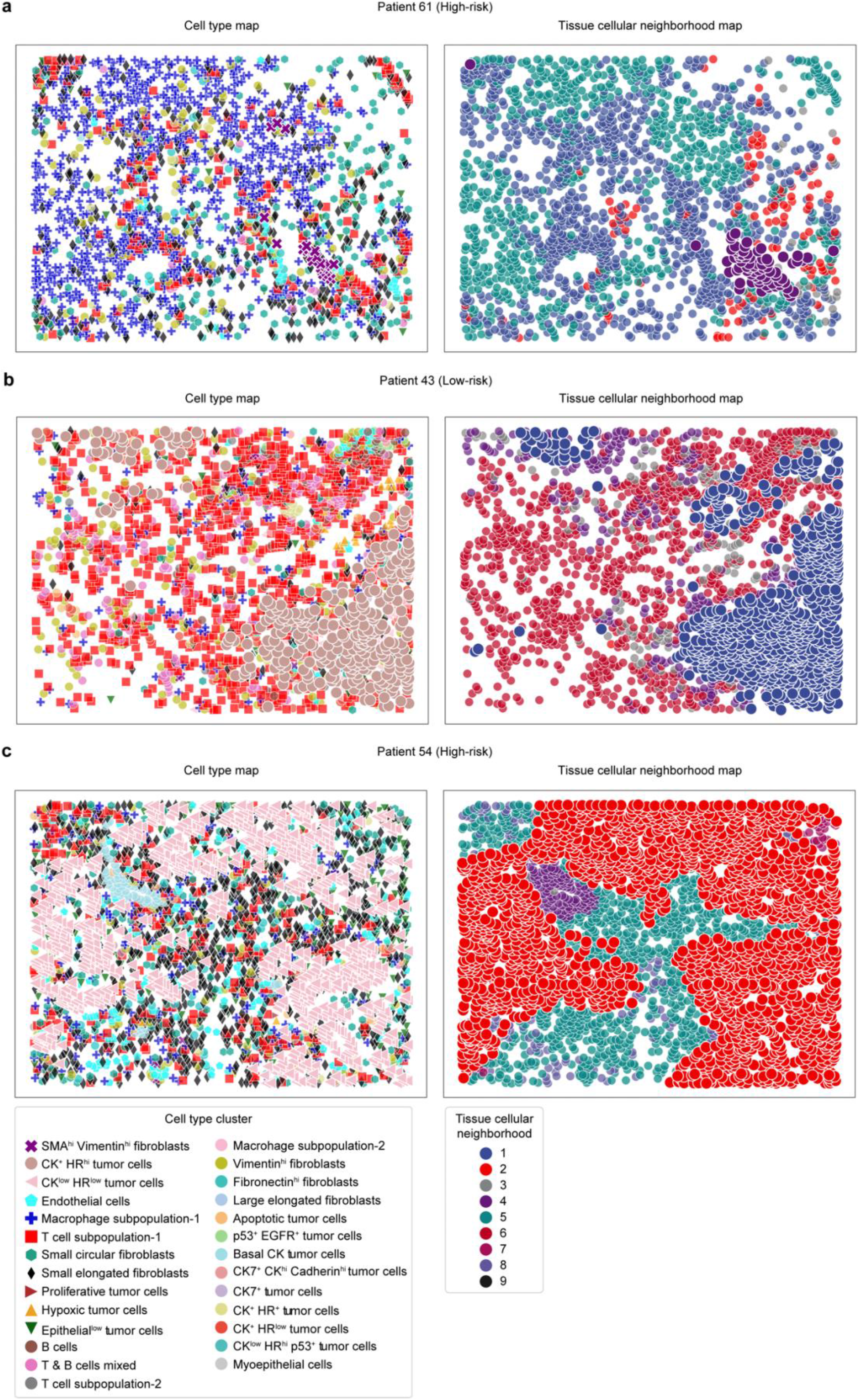
Representative tissue cellular neighborhoods specific to low- and high-risk breast cancer patients. **(a)** Cell type and tissue cellular neighborhood (TCN) maps of a representative high-risk patient with SMA^hi^ Vimentin^hi^ fibroblasts enriched in TCN-4. **(b)** Cell type and TCN maps of a representative low-risk patient with CK^+^ HR^hi^ tumor cells enriched in TCN-1. **(c)** Cell type and TCN maps of a representative high-risk patient with CK^low^ HR^low^ tumor cells enriched in TCN-2. For clarity, cells of studied types and TCNs are shown in large size without transparency in all cell type and TCN maps.

**Supplementary Figure 5.**
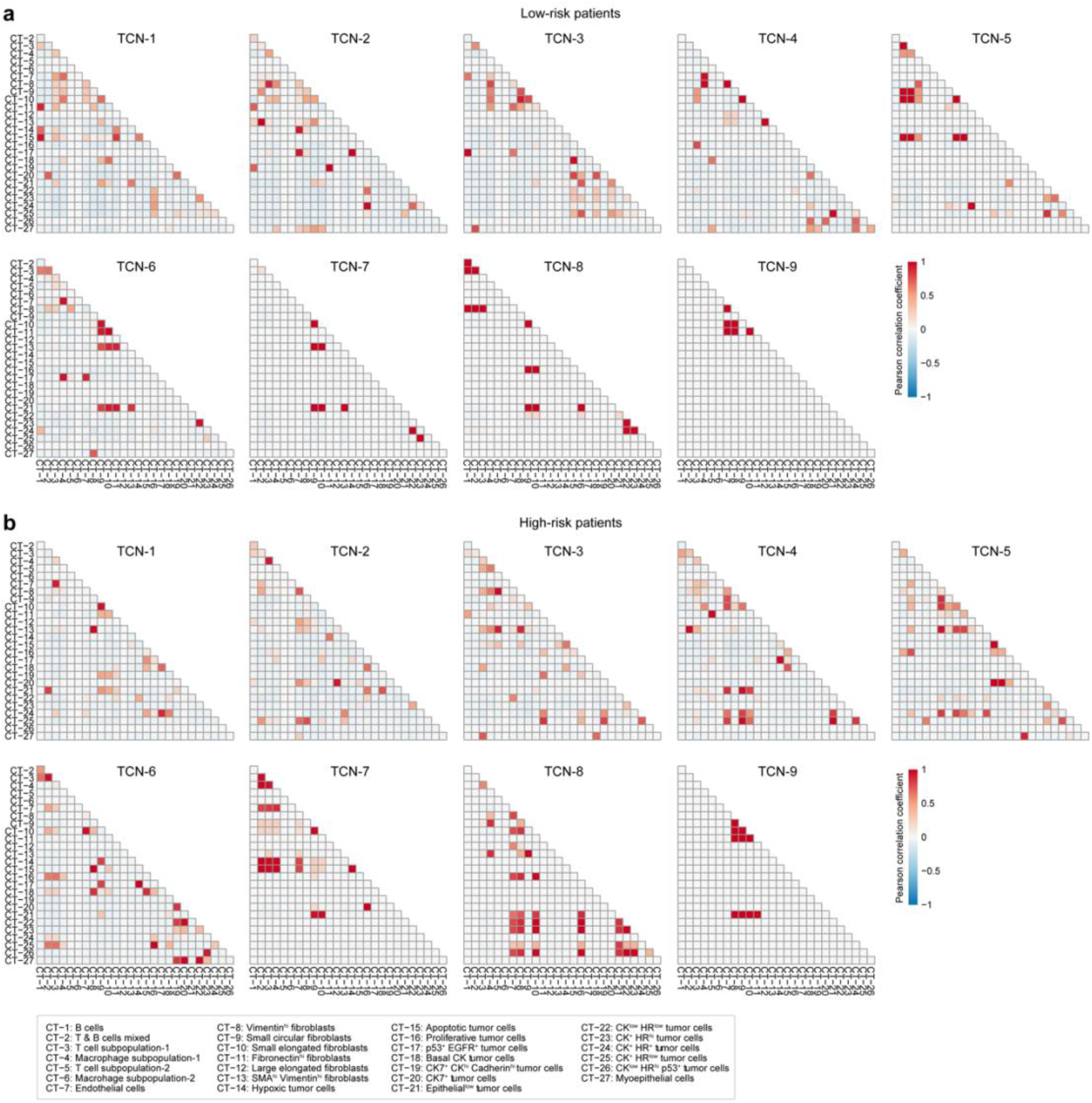
Cell type associations within tissue cellular neighborhoods in breast cancer. Heatmaps for correlations of the enrichment scores of any two cell types (CTs) within each of the nine tissue cellular neighborhoods (TCNs) identified in low-risk **(a)** and high-risk **(b)** patients.

**Supplementary Figure 6.**
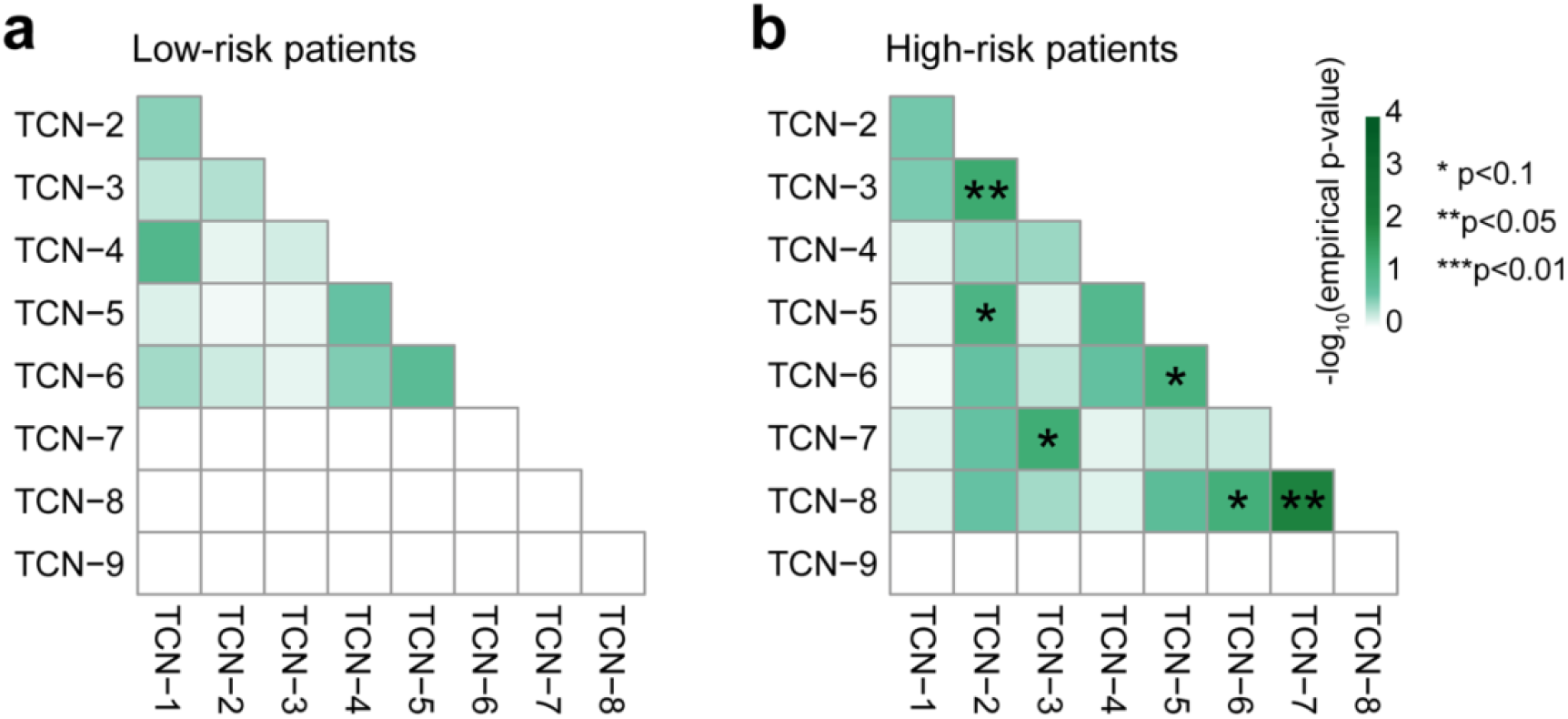
Cell type associations between tissue cellular neighborhoods in breast cancer. Heatmaps for empirical p-values of canonical correlation coefficients of the cell type enrichment scores of tissue cellular neighborhood (TCN) pairs in low-risk **(a)** and high-risk **(b)** patient groups. P-values were computed using permutation test-based *p.perm* function from the R package “CCP (v1.2)”.

**Supplementary Figure 7.**
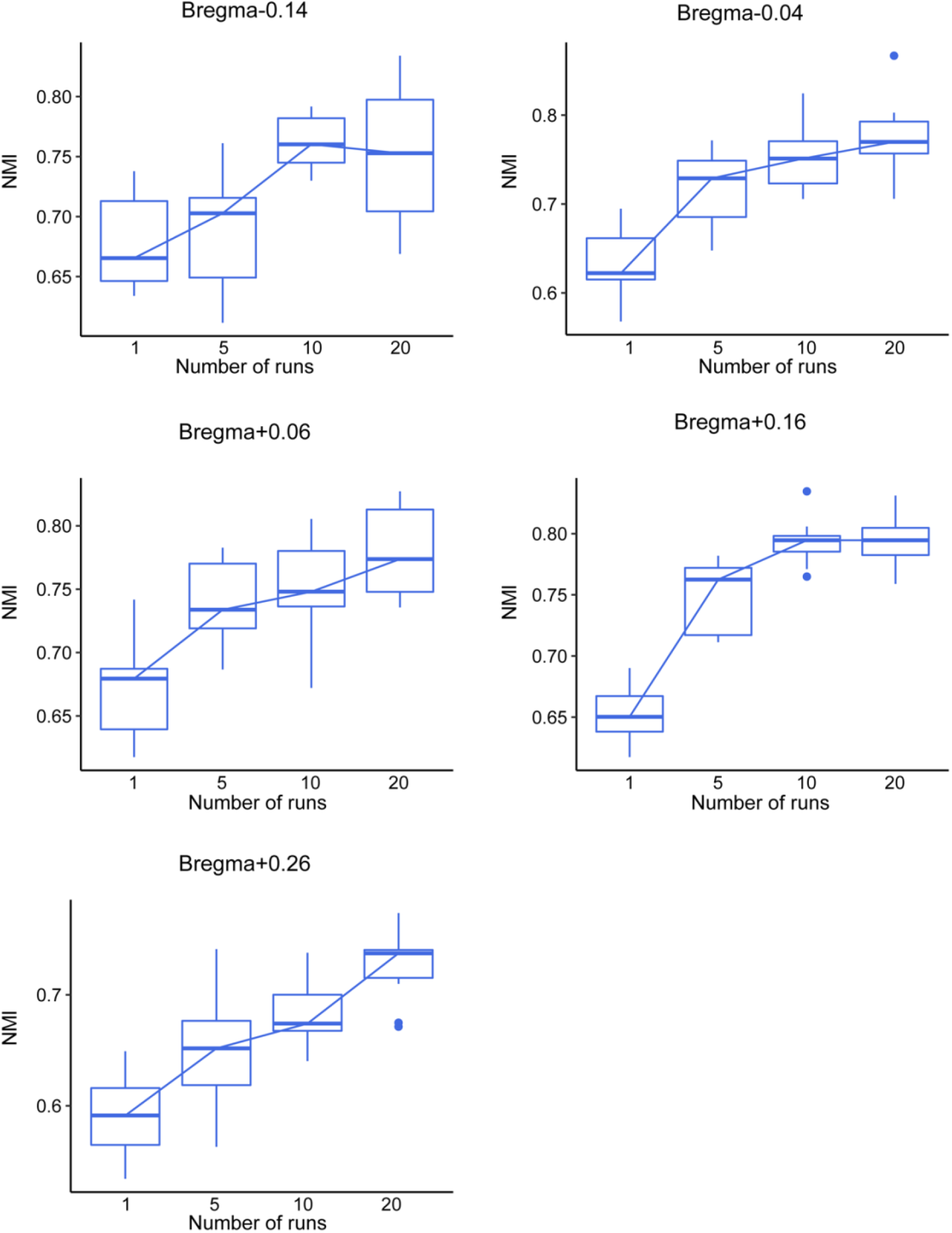
Robustness of CytoCommunity on the MERFISH dataset. CytoCommunity was applied to each MERFISH image for five times. Each time, different numbers of runs of the soft tissue cellular neighborhood (TCN) assignment module was conducted. Normalized mutual information (NMI) between any two TCN partitions generated by the five sets of experiments are shown in boxplots with lines connecting the median of each group.

